# Modeling decision-making under uncertainty: a direct comparison study between human and mouse gambling data

**DOI:** 10.1101/570499

**Authors:** Lidia Cabeza, Julie Giustiniani, Thibault Chabin, Bahrie Ramadan, Coralie Joucla, Magali Nicolier, Lionel Pazart, Emmanuel Haffen, Dominique Fellmann, Damien Gabriel, Yvan Peterschmitt

**Author notes:** Corresponding author Peterschmitt Y.

## Abstract

Decision-making is a conserved evolutionary process enabling to choose one option among several alternatives, and relying on reward and cognitive control systems. The Iowa Gambling Task allows to assess human decision-making under uncertainty by presenting four cards decks with various cost-benefit probabilities. Participants seek to maximize their monetary gains by developing long-term optimal choice strategies. Animal versions have been adapted with nutritional rewards but interspecies data comparisons are still scarce. Our study directly compared physiological decision-making performances between humans and wild-type C57BL/6 mice. Human subjects fulfilled an electronic Iowa Gambling Task version while mice performed a maze-based adaptation with four arms baited in a probabilistic way. Our data show closely matching performances among species with similar patterns of choice behaviors. Moreover, both populations clustered into good, intermediate, and poor decision-making categories with similar proportions. Remarkably, mice good decision-makers behaved as humans of the same category, but slight differences among species have been evidenced for the other two subpopulations. Overall, our direct comparative study confirms the good face validity of the rodent gambling task. Extended behavioral characterization and pathological animal models should help strengthen its construct validity and disentangle determinants of decision-making in animals and humans.

## Introduction

Do animals gamble for food just like humans gamble for money? Most species, including human beings, have to make accurate assessments of cost-benefit probabilities in reward-driven contexts. For example, the longer an animal invests searching for food, the most accurate predictions will be made about which sources are the most rewarding. Meanwhile though, the probability to experience an adverse event, like meeting a predator, will also increase. Decision-making (DM) is thus an essential mechanism for survival in partly predictable situations^1^. The underlying principles of how living beings achieve efficient DM are still not fully understood, and assessing human DM by using animal models requires yet further validation. As for humans, laboratory tests simulating real-life DM have been developed, such as gambling tasks. The Iowa Gambling Task (IGT) in particular is widely used to assess human DM under uncertainty. The optimal choice strategy requires maximizing monetary gains by selecting cards from four decks with various cost-benefit probabilities^2^.Players face a choice conflict between playing from long-term disadvantageous decks yielding higher gains but frequent losses, or from advantageous ones associated with smaller immediate rewards and less frequent penalties. Thus, optimal decisions consist in refraining preference for larger immediate rewards favoring long-term benefits from lower but more frequent gains.

The IGT was originally created to study DM impairments in patients with ventromedial prefrontal cortex damage. Compared to healthy participants who progressively learn to select the advantageous decks, these patients perform defectively, being unable to set the optimal strategy from repeated negative outcomes. Besides, many IGT studies have shown a high interindividual variability regarding the performances in a healthy population^3^-^5^. Indeed, several clinical reports indicated that, whereas a majority of subjects develop the optimal strategy, others do not acquire a preference for one deck over the others, which is indicative of a lack of learning^3,6,7^. Moreover, up to one third of healthy subjects keep on choosing the disadvantageous decks^5^. These data illustrate a behavioral continuum with overlap in choice strategies between human physiological and pathological conditions.

Interestingly, animal versions of the IGT have been developed, with species appropriate adjustments, to assess DM in a design comparable to humans^8^-^16^. For example, in rodent gambling tasks the decks of cards have been replaced by mazes or operant chambers suited with different options of varying outcomes. Besides, monetary gains become nutritional rewards since money and food are thought to drive similar behaviors^17^.

Similar to human studies, lesioned rodents perform suboptimally in adapted IGT versions, as well as they show delayed DM compared to healthy individuals^18,19^. For the majority of healthy rodents, an improvement of performances is observed as the task progresses and individuals commonly cluster in three subpopulations reflecting variable choice strategies^12,14,20^. Good DM individuals quickly develop a strong preference for the advantageous options, while poor DM individuals display the worst performances, either not showing any preference for neither advantageous nor disadvantageous options, or displaying a long-term preference for the disadvantageous ones. Intermediate DM individuals perform between the other two subgroups. Whether these interindividual variabilities observed in both healthy human and animal populations are similar has never been directly assessed to further substantiate face validity of models.

Literature has shown that many factors may account for good and poor DM performance. For example, Bechara et al. (2000)^21^ explain poor IGT performance in terms of atypical sensitivity to reward or punishment. Other studies show that adopting a rigid behavior, i.e. selecting persistently the same option without switching between the available ones, or a flexible behavior with sustained exploration of every option, were also indicative of endpoint performances^22,23^. Furthermore, animal and human performances in gambling tasks are usually measured within distinct studies, being therefore indirectly compared, and even if animal gambling tasks have been specially created to simulate human DM processes. Comparative research has indeed shown how challenging comparisons among species can be^20^.

Here, we aimed to bridge findings from clinical and preclinical research on DM processes featuring uncertainty. To this purpose, we directly compared human and mouse physiological performances, for the first time in the same study, by using IGT-adapted tasks and common analytical methods.

Based upon recent literature addressing mouse gambling tasks (mGT) validity^12,16,19,20,24^, we forecasted a good face validity of our animal model with similar overall performances regarding the human population^25^. To check for the existence of similar variability in choice strategies among species, we then compared the stratification of both populations according to endpoint performances. To evaluate the construct validity, we further studied behavioral indexes of cognitive processes subserving optimal choices. Finally, we sought for correlations with endpoint performances, focusing on parameters of rigidity^26^, flexibility^20^ and win-stay and lose-shift choices^27^. We also analyzed psychometric scores from human subjects and reward sensitivity data in mice, to relate performances or choice strategies to additional behavioral traits.

## Results

### Gambling performances in human and mice populations (Figure 1)

In order to circumvent variations in the designs of human and mouse gambling tasks and to allow for direct comparison of data, we analyzed performances in terms of advantageous choices as a function of task progression, expressed in percentage. Therefore, total number of trials (see Methods section for details) was split into five 20% of trials-blocks.

**Figure 1.**
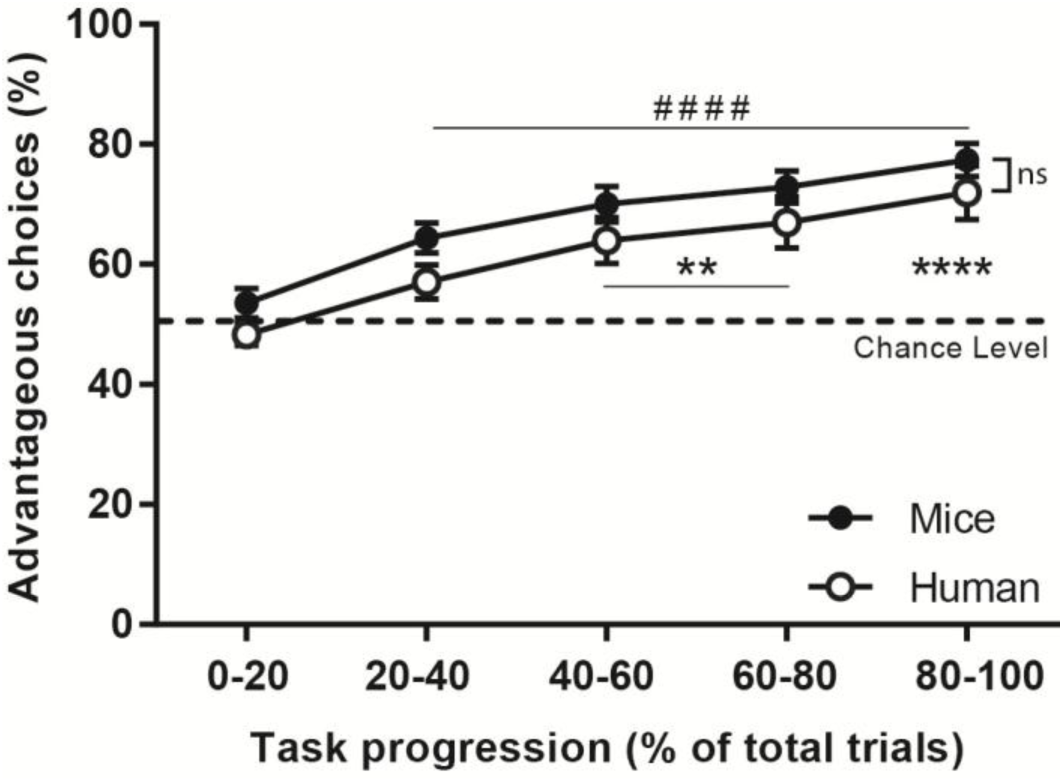
Gambling performances in human and mice populations. Performances expressed as percentage of advantageous choices (mean ± SEM) during task progression in blocks of 20% of total number of trials reveal similarities among species. Comparison by repeated measures ANOVA shows no significant (ns) differences between species. Human performances differ from chance level from the third 20%-block onwards, while mouse performances are already different form the second block (t-test: advantageous choices different from chance level, mice: ^*####*^, p<0.0001; human: **, p<0.01: ****, p<0.0001).

Our IGT data show that humans acquired task contingencies after completion of 40 % of the task, performing above chance level from the third 20%-block onwards (Student’s test (t-test), p<0.01). A learning effect has been evidenced as performances gradually improved over time (F(4,156) = 15.4; p<0.0001) after 20% of the task was completed and till the end (p<0.0001, above behavioral output of the 1^st^ 20%-block).

As regard the mGT data, mice performed above chance level after the first 20% of the task, thus oriented towards favorable choices earlier than humans did (t-test, p<0.0001). Performances progressively improved over time (F(4,195) = 11.9; p<0.0001) already from the second 20%-block (2^nd^ block: p<0.05, 3^rd^ block: p<0.001, 4^th^ and 5^th^ blocks: p<0.0001) and compared to the beginning of the task.

The comparison of IGT and mGT performances show no significant effect of species in the percentage of advantageous choices (F(1,78) = 3.1; p=0.80). Both populations performed also similarly along task progression, as shown by the absence of statistical difference in the interaction between factors (species and 20%-Blocks) (F(4,312) = 0.06; p=0.99).

These results show that both species learn the task contingencies and indicate that mice and humans display a similar time course of performances’ improvement with curves perfectly matching.

### Comparable categories of good, intermediate and poor decision-makers (DMs) with equal proportions among species (Figure 2)

Individual data have been plotted showing larger dispersion in humans than in mice (Fig. 2, A1 and B1). Intraspecies variability has been investigated by k-mean clustering stratification according to endpoint performances, discriminating three subpopulations with different choice strategies: good, intermediate and poor DMs (see Methods section for details). Individuals overtly displaying the optimal strategy were referred as good DMs and represent 42.5% of the human population (mean percentage of advantageous choices ± SEM: 97.4 ± 0.8) and 40% of the mice population (91.3 ± 1.5). Poor DMs remained around 50% of advantageous choices, with no significant preference neither for advantageous nor for disadvantageous options, and represent 25% of humans (34.3 ± 5.2) and 22.5% of mice (54.1 ± 2.5). The third subgroup corresponds to individuals that developed a preference towards some options, although they did not find the most favorable strategy. These intermediate DMs represent 32.5% of humans (64.5 ± 2.4) and 37.5% of mice (73.8 ± 1.0). The proportions of each subpopulation in humans and mice were compared and no significant differences were found (Chi-Square test = 0.011, p=0.99) (Fig. 2, A2 and B2).

**Figure 2.**
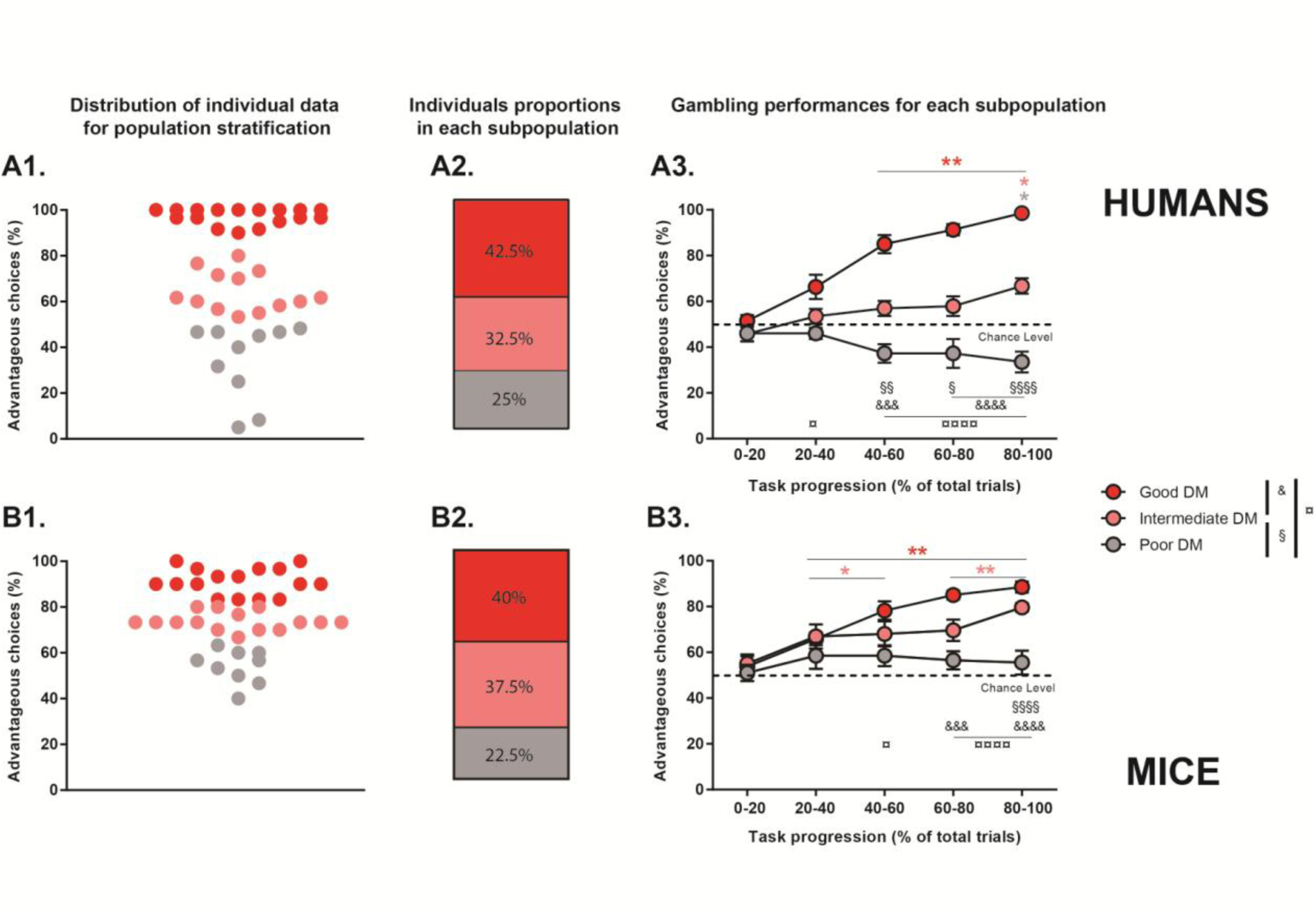
Comparable categories of good, intermediate and poor DMs with equal proportions among species. Distribution in humans (A1) and mice (B1) of individual percentage of advantageous choices at the end of the task used for k-mean clustering stratification. Proportions in humans (A2) and mice (B2) of individuals in good (red), intermediate (pink) and poor (grey) DM subpopulations. Gambling performances in human (A3) and mice (B3) subpopulations expressed as percentage of advantageous choices (mean ± SEM) during task progression in blocks of 20% of total number of trials. W test to show group performances different from chance level (advantageous choices different from 50%: *, p<0.05; **, p<0.01; ***, p<0.001). MW tests to further show group differences (good versus intermediate: &, p<0.05; &&, p<0.01; &&&, p<0.001; &&&&, p<0.0001; good versus poor: ¤; intermediate versus poor: §).

In humans, good DMs performed above chance level after 40% of the task was completed (Wilcoxon (W), p<0.01), while intermediate and poor DMs differed from it only in the last 20%-block (p<0.05). Furthermore, the ANOVA analysis revealed a significant interaction between Clusters and 20%-blocks (F(8,148) = 16.2; p<0.0001). Good DMs performed differently than poor DMs from the second 20%-block (Mann Whitney (MW), 2^nd^ block: p<0.05, 3^rd^ to 5^th^ blocks: p<0.0001), and differently than intermediate DMs in the last 60% of the task (3^rd^ block: p<0.001, 4^th^ and 5^th^ blocks: p<0.0001). Intermediate and poor DMs performed differently from the third block onwards (3^rd^ block: p<0.01, 4^th^ block: p<0.05, 5^th^ block: p<0.0001) (Fig. 2, A3).

Regarding mouse data, intermediate and good DMs performed above chance level early after 20% of the task was completed (W, good 2^nd^ to 5^th^ blocks: p<0.01; intermediate 2^nd^ and 3^rd^ blocks: p<0.05, 4^th^ and 5^th^ blocks: p<0.01), while poor DMs never differed from chance level (p>0.05). The ANOVA analysis also revealed a significant interaction between factors (F(8,185) = 5.3; p<0.0001). Mice good DMs performed differently than poor DMs after 40% of the task was completed (MW, 3^rd^ block: p<0.05, 4^th^ and 5^th^ blocks: p<0.0001), and differently from intermediate DMs later on (4^th^ block: p<0.001; 5^th^ block; p<0.0001). Mice intermediate and poor DMs performed differently only during the last 20% of the task (p<0.0001).

When comparing gambling performances in mice and humans for each subpopulation, differences were found for intermediate DMs (MW, p<0.05), with mice making more advantageous choices than humans. On the contrary, humans good DMs achieved more advantageous choices than mice of the same subgroup (p<0.01). Furthermore, humans poor DMs made worse decisions than mice (p<0.01).

Collectively these data reveal that both mice and human populations clustered into three comparable DM categories, with closely matching proportions and displaying similar performances, though with more extreme choices in the human population.

### Relationship between DM performances and choice behaviors in humans and mice (Figure 3 and Supplementary data)

Maximization of benefits and reduction of costs characterizing optimal performances require flexibly adapting to contingencies in order to favorably orient future choices. To compare DM strategies, behavioral measures of cognitive processes at the beginning and the end of the task have been calculated and correlated with endpoint performances (see Methods section for details).

**Figure 3.**
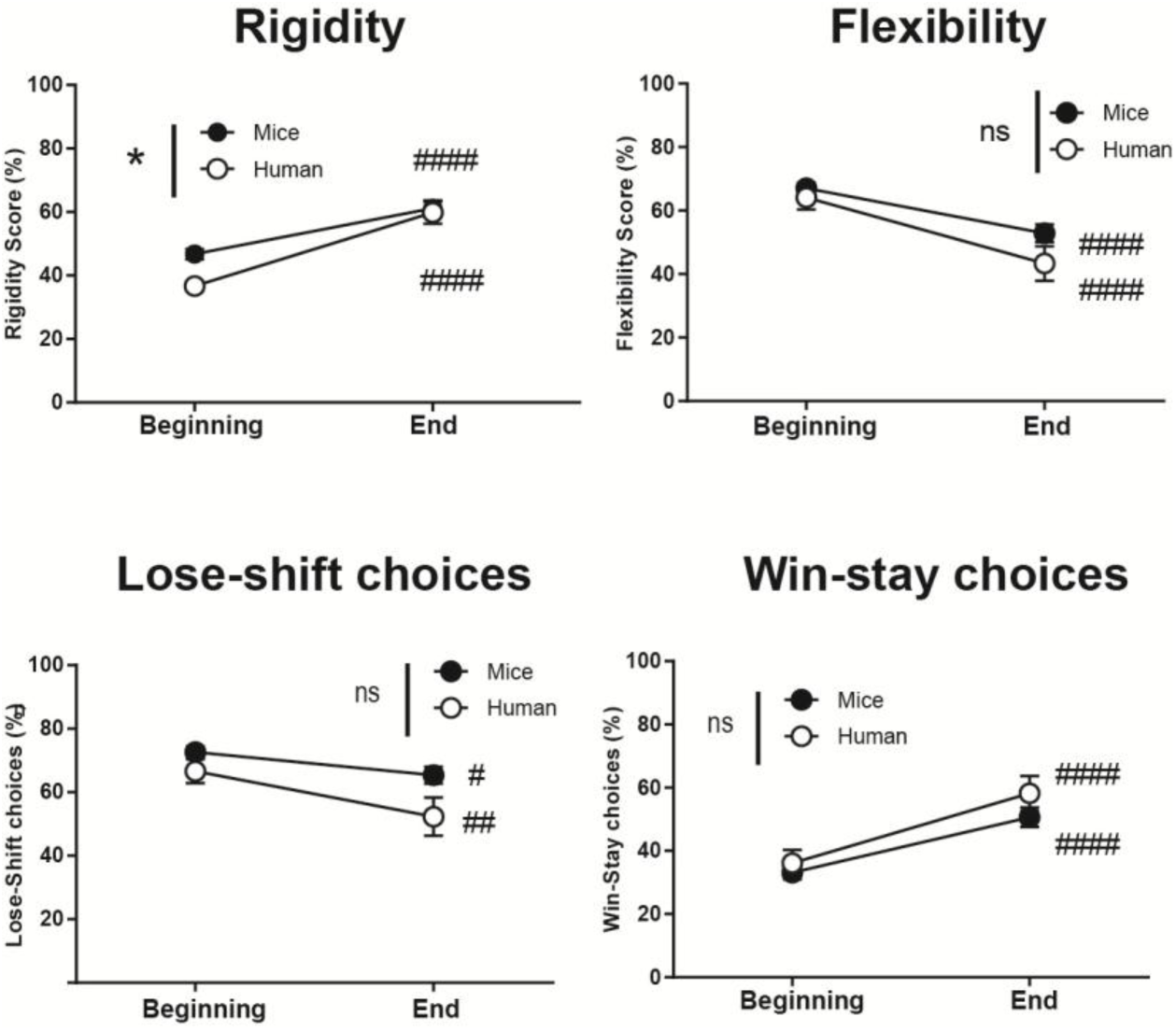
Evolution of choice behaviors during task progression. Significant progression of the rigidity, flexibility, lose-shift and win-stay choices for human (white circles) and mice (black circles) populations, between the beginning and the end of the experiment. Comparison of choice strategies at the population level by repeated measures ANOVA and post hoc t-tests (beginning versus end of the task: #, p<0.05; ##, p<0.01; ####, p<0.0001; humans versus mice: *, p<0.05).

#### Correlations between endpoint performances and behavioral determinants of DM (Supplementary data Figure 1)

In humans, rigidity scores significantly correlated with final performances at the beginning (r=0.428, p<0.05) and at the end of the task (r=0.489, p<0.01). Flexibility scores showed a negative correlation at the beginning of the task (r=-0.509, p<0.01), which disappeared at the end (p>0.05). No significant correlation was found for lose-shift choices independently of the moment of the experiment. However, win-stay choices at the beginning of the task significantly correlated with final performances (r=0.518, p<0.01).

In mice, like in humans, rigidity scores significantly correlated with endpoint performances at the end of the task (r=0.669, p<0.001), but not at the beginning (p>0.05). On the contrary, no correlation was found for flexibility at the beginning of the task (p>0.05), but a negative correlation at the end (r=-0.595, p<0.001). In the same way as humans, no correlation was found regarding lose-shift choices (p>0.05), but a significant correlation between win-stay choices and final performances at the end of the task (r=0.582, p<0.001).

In brief, DM strategies in humans and mice rely on similar adaptive choice behaviors that correlate with endpoint performances.

#### <u>Evolution of choice behaviors during task progression (Figure 3 and Supplementary data Figure</u> <u>2)</u>

In both humans and mice a significant effect of time course was found for flexibility (F(1,78) = 48.4, p<0.0001), lose-shift (F(1,78) = 15.8; p<0.001) and win-stay scores (F(1,78) = 51.9, p<0.0001). (Fig. 3).

Whereas no global difference among species was found for any parameters, a significant interaction between factors (species and time course) was found for rigidity (F(1,78) = 4.6, p<0.05). If humans and mice selected more often the same options along the task (t-test, p <0.0001), humans were significantly less rigid than mice at the beginning of the experiment (p<0.05), whereas no difference was observed at the end (p=1) (Fig. 3).

The interspecies comparisons revealed that, at the beginning of the task, humans intermediate and poor DMs were significantly less rigid than corresponding mice (MW, intermediate: p<0.0001; poor: p<0.01), but not good DMs (p>0.05). At the end of the task, only humans intermediate DMs were still less rigid than mice (p<0.01). Flexibility scores were similar in both species for intermediate and poor DMs at the beginning of the task (p>0.05), but humans good DMs were less flexible than mice of the same group (p<0.05). At the end of the experiment, humans good DMs continued being less flexible than mice (p<0.001) and no differences were found between the other two groups (p>0.05). No subgroup differences in lose-shift and win-stay choices were observed between populations at the beginning of the task (p>0.05). At the end, only mice good DMs were more prone to switch from option after a penalty than humans of the same subgroup (p<0.05). However, humans good DMs continued choosing the same option after a positive outcome more frequently than mice (p<0.01). No differences were found for the other subgroups (p>0.05) (see Supplementary data Fig. 2).

Overall, these data show that human and mice do not differ in their choice strategies at the population level, but suggest slight differences in intermediate and poor DMs subpopulations among species. Remarkably, mice and humans good DMs behaved alike.

## Discussion

The goal of this study was to directly compare DM under uncertainty between humans and mice using IGT adaptations according to the litterature^9,12,25^. To reduce conceptual and methodological differences between the tasks, we controlled factors known to interfere with the results such as sex differences and the presence of instructions, especially in humans. In that respect we recruited only male subjects because they have been described as less risky than females, choosing the advantageous options more frequently in the IGT^28^ and its rodent adaptations^9^. Our study shows perfectly matching performance curves as a result of comparing human and mice data as a function of task progression. Thus, mice performing 100 multi-session-trials reached equivalent endpoint performances as humans completing 200 trials within a single session. Our results demonstrate that mice’s behavior when they are rewarded with food closely resembles to that observed in humans who are rewarded with money, both populations being able to discern advantageous options in the long-term among the other possibilities. Humans and mice started with an explorative search, displaying equal preference for either the advantageous or disadvantageous options. A preference for advantageous choices progressively emerged during what has been referred to as the exploitation phase^8,15,29^. If overall performances were very similar, mice however more promptly selected advantageous options than humans, requiring less exploratory trials to adopt the favorable strategy. The learning curves observed in our mice confirm previous animal results obtained in similar conditions^12,26^ and in other variants of rodent gambling tasks^20,27^. These findings are also in agreement with human literature reporting on the one hand, longer exploration phase when instructions about the presence of advantageous and disadvantageous decks are not provided, and on the other hand, an improvement of the endpoint performances in direct relation to the total number of trials^30^-^32^. Since our participants did not receive task instructions, they required longer in figuring out the strategy to elaborate^32,33^. Similarly, the emergence of the preference for an option has been shown to appear earlier when rodents are taught about the gambling task contingencies^14^.

The alignment of performances in humans and mice using either money or food as reward is essential because the nature of reward is considered a major limitation for the animal versions of the IGT^14,26^. Modeling loss of reward in animals in a similar manner as in humans is a challenge. Food aids in the survival of species: it is a primary reinforcer, since it strengthens behavior and satiates the basic biological drives. Money is a secondary reinforcer: its value is relative to the primary reinforcer. Hunger and satiety are factors difficult to control, which patently influence animals motivational state^34,35^. However, since the interest for money is also difficult to control due to its subjective nature, it raises similar concerns and can lead to the same consequences. Although other animal studies have proposed delays and absence of reward as penalties^8,10,14,36^, the development of new tasks to overcome this issue is also needed.

In addition to the challenge associated to reward nature and processing, the internal state (see *the somatic marker theory*^37^) and the context also generate differences in behavior. Humans play at a computerized version of a game, whereas mice have to physically explore a maze to get the rewards. In that respect, new automated touchscreen gambling protocols have been developed for animals and humans^38^. Animal automated testing in operant chambers would also help deepen analysis of motivational aspects for instance.

If similar overall gambling performances underline proper face validity of our mouse model, construct validity should be substantiated by similar choice strategies in humans and mice. Those can be investigated from additional cognitive proxies subserving behavior^19^. A closer look at the endpoint performances revealed that they correlated with similar choice behaviors in both populations, whose relative contributions either took over or faded away from initiation to completion of the task. In fact, as advantageous choices increased along trials progression, preference for one option progressively emerged (increased rigidity), while less options were explored (decreased flexibility), with individuals becoming more sensitive to reward (increased win-stay choices) and tending to more easily cope with penalties (decreased lose-shift choices).

Our results also highlighted close common interindividual variability in mice and human populations, when clustered into three subgroups of individuals, those exhibiting different behavioral strategies. These subgroups, ranging from good, to intermediate and poor DMs, perfectly match their proportions between species. Additionally, their final performances correlated with choice behaviors, supporting the assumption that DM outcomes in humans and mice rely on comparable choice strategies and that our animal preparation and more generally the rodent model of gambling present good face and construct validities.

Concerning endpoint performances, interindividual variabilities show a larger spreading in the human population, accounting for better and worse decisions. The upper extreme scores in humans might be a consequence of a longer gambling design. Besides, inbred animals such as C57BL/6J used in the present study, which are known to behave uniformly and display low interindividual variability^39^ (but see recent Tuttle et al.’s work^40^), were food restricted, which might motivate exploration and the completion of the nutritionally rewarding task^14^. That could also explain why our mice never chose preferably the disadvantageous options. Noticeably, very low performances in poor DMs animals more reminiscent to human data than ours, have been published^14,15^, illustrating that performance’s output highly depend on task design.

Further, the stratification analysis revealed salient matching proportions between human and mice subpopulations, strengthening the face validity of our animal model. Individuals attaining the best performances –good DMs-, composed the largest subgroup in both species. Several human studies correlate predominantly good performances with the development of an optimal strategy, neglecting aversion to risk^41,42^, as in mice studies where good performances are related to a secure strategy^12^. The design of the task does not allow to firmly establish whether participants develop an explicit knowledge or display some aversion to risk. Presumably, in our study both situations were present in some individuals. Interestingly, mice good DMs needed fewer trials than humans to perform above chance level. During the exploitation phase, humans good DMs developed stronger preference for one option (increased rigidity) over intermediate and poor DMs, whereas mice good DMs differed only from poor DMs. The evolution of penalty aversion was also similar for both species, with good DMs showing significantly less lose-shift choices than intermediate DMs. These results suggest that cognitive strategies underlying DM performances in both species might be similar, at least for the good DM subgroup.

Individuals conforming the second subgroup –intermediate DMs-, while selecting more often advantageous choices, maintained a high level of exploration of all options (constant flexibility), without neglecting the disadvantageous choices. These intermediate DMs are slower learners than good DMs and did not achieve the best strategy to maximize their rewards. However, IGT studies have shown that these performances can be significantly enhanced with additional trials^43^. Besides, an earlier emergence of the exploitation phase has been shown in mice compared to humans. Furthermore, mice intermediate DMs displayed an increase in win-stay choices along the task progression whereas humans did not, suggesting that intermediate categories might not completely overlap between species.

The subgroup with the worst performances –poor DMs-maintained the exploration of all available options, even if associated to uncertain outcomes. Individuals of this subgroup did not manage to find a favorable strategy or to develop any implicit knowledge of the task. However, humans poor DMs ended the task performing significantly below chance level in terms of advantageous choices, while that did never happen in the animal population. Poor DMs of both populations exhibited a high flexibility, suggesting an ineffective exploration of the available options. Mice from this subgroup have been proposed as models for vulnerability of pathological gambling or addiction^12,24^, in the same line of human studies seeking for behavioral markers of pathological predisposition or endophenotypes^44,45^. Indeed, these animals seem less risk-averse, a trait that has been already interpreted as an indicator of weaker cognitive control over immediate loss^27^. Nevertheless and contrary to mice, some humans poor DMs developed a real preference for disadvantageous options. This kind of deleterious preference however, has been described in mice following singled-session mGT protocols^20^ and other rodent IGT versions^14,15^.

Similar overall performances and comparable individual gambling strategies observed between both species suggest that they share conserved cognitive processes essential for successful DM. Several cognitive functions related to attention, discrimination and reversal learning are necessary to perform optimally at the IGT. Good performances arise from planning flexibility, monitoring incoming information, evaluating risk-reward and refraining from choosing short-term advantageous options. In that respect, a wide range of comparative approaches have been proposed to discern between cognitive processes, in rodents and in humans, which are activated when performing cognitive tasks, both at the attentional and the mnemonic levels^46^. Literature has also shown that there is a strikingly similar range of cognitive abilities between rodents and humans, as well as a remarkably high degree of anatomical overlap in their brain functions^47^. Rodents are even able to outperform humans in some learning tasks^48^ but we do believe that our slight differing kinetics are mainly accounted by task design variability. Rodents possess preserved metacognitive abilities, which is an essential aspect of the IGT^49^. Indeed, patients suffering from metacognitive deficits such as those with substance use disorders, show poorer performances at the IGT^4^. Pathological gamblers not only perform poorly at the IGT, but also erroneously estimate that their performances are much better than they actually are, which is referred to as subjective biases^21^. Whether subjective biases also determine choices in mice poor DMs remains to be evaluated.

Although different behavioral aspects of DM have been assessed in order to better interpret the identified strategies, a general common pattern between humans and mice has not been fully revealed. By themselves, rigidity, flexibility and sensitivity to positive and negative outcomes cannot explain the evolution of performances, neither their emergence, in the three subgroups equally. The IGT alone unfortunately does not allow distinguishing reward maximization from ambiguity aversion for instance, being its output insufficient to determine why a subject selected an option. In this perspective, further behavioral characterization has been attempted. In humans, we evaluated by the behavioral inhibition system (BIS) and the behavioral activation system (BAS) scales, the motivation to avoid aversive outcomes and to approach goal-oriented outcomes respectively (see Supplementary data). Endpoint performances did not correlate with neither BIS/BAS scores in humans (data not shown). In parallel, reward sensitivity assessed by the sucrose preference task in mice (see Supplementary data), did not significantly differed between subgroups. These results contrast with those from Granon’s team, where a stronger sucrose preference is described for good (“safe”) compared to poor (“risky”) mice DMs^12^. Reward sensitivity did not correlate with endpoint performances neither (data not shown). Conversely, we did observe a correlation between sucrose preference scores and lose-shift choices at the conclusion of the task (data not shown). These apparent discrepancies could be accounted for by protocol variations and suggest complex relationships between reward sensitivity and DM strategies. This is in line with former suggestions about poor performance being likely mediated by sensitivity to high reward^20^. The parameters we evaluated in both species are insufficient to draw a firm conclusion for the relation between reward maximization and risk aversion. These behavioral traits cannot solely explain the differences on the acquisition of the general contingencies of the task during the exploration phase, nor alone account for endpoint performances. Further extensive behavioral characterizations are required in both populations to better understand determinants of DM and relationships between reward and cognitive control systems.

## Conclusion

Food, like money, influence DM and risk-taking behavior^35^.The data thus far suggest that rodents behave in a way similar to humans, that is, they tend to choose the option with the best long-term payoff more often as the task progresses. Our results point to similar patterns of choice behaviors present across species. Accurate and validated animal models are indispensable crucial to study brain regions and circuits involved in DM. Identifying behavioral traits related to poor DM as pathological gambling endophenotypes, therefore requires experimental designs carefully controlling environmental conditions and genetic variations^8^. To conclude, our results directly support good face validity of the mouse version of IGT. The determinants of interspecies differences in choice strategies have to be explored in depth, although restricted variations observed seem so far insufficient to question the construct validity of the animal model. Future studies with extended behavioral characterization and pathological animal models should help disentangle processes subserving choice strategies, clarifying DM in animals and humans.

## Methods

### Participants

#### Humans

Forty healthy right-handed subjects, all male (mean age ± SEM = 24.7 ± 5.1; range 19-38), were involved in the study. None of them reported previous medical history of psychiatric disorders, substance or alcohol abuse, neurological diseases, traumatic brain injury or stroke, and none did report taking any medication.

Participants received information regarding the aim of the task and gave their written informed consent to take part in the study. Given the influence of real money playing a significant role on motivation, subjects were informed that the monetary payment will be proportional to the global gain obtained in the task^50^-^52^. Due to ethical considerations and whatever their performance, all participants received the maximum amount of 85€ at the end of the experiment. The protocol was approved by the Committee of Protection of Persons (CPP-Est-11 ; authorization given by the General Health Administration (ANSM 2016-A00870-51 and NCT 02862821)).

#### Mice

Forty male C57BL/6JRj mice (Ets Janvier Labs, Saint-Berthevin, France) were used for this study. All mice were 3-5 months old at the time of testing, were group-housed and maintained under a 12 hour-circadian cycle, with constant temperature (22 ± 2<u>°</u>C). Water was available *ad libitum* and all mice were food restricted at 80-90% of their free-feeding weight (mean weight (g) ± SEM = 22.2 ± 0.2). Experiments were performed in behavioral rooms, with tight luminous intensity. All procedures met the NIH guidelines for the care and use of laboratory animals and were approved by the University of Franche-Comte Animal Care and Use Committee (CEBEA-58). All efforts have been taken to minimize animal suffering during the testing according to the Directive from the European Council at 22^nd^ of September 2010 (2010/63/EU).

### Experimental procedure

#### Humans

The task was an adapted electronic version of the IGT^25^, whose aim was to win as much money as possible by making successive selections between four decks. Their composition, values and schedules reward-penalty were predetermined identically to the original form of the IGT^2,53,54^. Decks looked identical but they differed in composition. Decks A and B were disadvantageous: they yielded immediate rewards but in the long-run involved major economic losses. Decks C and D were advantageous: they yielded frequent small wins and smaller long-term penalties, which resulted in long-term gain. To adapt the IGT to our French population, the money used to play was converted from US Dollars to Euros. At the beginning of the task, participants had a loan of 2,000€.

Contrary to most IGT experiments, no specific instructions were given to participants regarding the presence of advantageous or disadvantageous decks, nor the number of trials, avoiding a somewhat partial advantage compared to animals^55^ (but see Rivalan’s work^18^). In absence of instructions, final performances usually worsen, the exploration phase therefore lengthens and the optimal strategy is hardly found in 100 trials only^7,32,33^. However, when allowed more trials, many individuals performing poorly in the first 100 trials are able to achieve good final performance^30,31^. To that purpose, the number of trials was increased from 100 to 200.

#### Mice

DM was evaluated using a mGT adapted from published protocoles^8,12^. The experiment took place in a 4-arm radial maze (identical and equidistant arms, 37cm long and 5,7cm wide), completely opaque, with a common central zone used as a start-point. Mice were rewarded with grain-based pellets (20mg Dustless Precision Pellets® Grain-Based Diet, PHYMEP s.a.r.L., Paris, France) or punished with grain-based pellets previously treated with quinine (180mM quinine hydrochloride, Sigma-Aldrich, Schnelldorf, Germany). Quinine pellets were poorly palatable but eatables.

Prior to every experimental session, animals were acclimatized to the behavioral room during 30 minutes. The experimental design was composed of 5 blocks of 20 trials, over five days (a total of 100 trials per animal). The first 10 trials of each block took place during the morning, and the second 10 trials during the afternoon. Before the first trial of the first block, mice had 3 minutes to explore and eat inside the maze (*first habituation period*). From the second block, mice had 2 minutes to explore the maze before the first trial, but no food was available (*general habituation period*). After the respective habituation period, mice were placed at the start-point, inside an opaque cylindrical structure to avoid early orientation through future choices. The cylinder was removed after 5 seconds and animals were allowed to choose an arm. Mice had one minute to choose an arm, explore it and eat the reward. If the choice was not made in time, an extra-minute was given.

Our mGT has been adapted in order to minimize the effect of satiety during the task. For that, two arms gave access to a small reward (1 pellet) in the first trial of each half block (trials 1, 11, 21, 31, 41, 51, 61, 71, 81 and 91), and bigger rewards (3-4 pellets) in the other 18 choices of each block, with a small probability of presenting a punishment (3-4 quinine pellets, twice in 18 possible choices) (*advantageous arms*). The other two arms (*disadvantageous arms*) gave access to a bigger reward (2 pellets) in the first trial of each half block, but bigger punishments (4-5 quinine pellets) in the other 18 choices of the blocks, with a small probability of presenting a reward (4-5 pellets, once in 18 possible choices). Between consecutive trials, animals were replaced in their home cages during 90 seconds. The localization of advantageous and disadvantageous arms was randomized and the probability combinations were different for each animal.

### Determination of interindividual differences

A clustering method already used in mice gambling tasks^12,56^ was applied to look for interindividual differences in both human and mice populations. The optimum objects’ partition into a specific number of clusters was thus automatically found. This procedure minimizes the within-cluster variance and maximize the between-cluster variance^57^. In accordance with the literature^12^, the mean percentage of advantageous choices was calculated for the last 30% of the tasks, when performances were highly stable (p<0.01 in mice and humans). The individual performances were then divided in three groups (*good, intermediate* and *poor DMs*).

### *Choice behaviors: rigidity*, *flexibility*, *lose-shift and win-stay scores*

We measured the rigidity score of humans and mice by calculating the highest percentage of choice of a deck or arm. We determined the flexibility score by calculating the proportion of switches from one deck or arm to another. The lose-shift score, as a measure of negative outcome aversion, was assessed by calculating the proportion of switches after a loss of money or a quinine penalty outcome. In the same line, win-stay scores were calculated as the proportion of remaining with the same option after receiving a reward. Each behavioral measure was calculated at the beginning (first 40% of the task) and at the end of the experiment (last 40% of the task), according to previous studies^12^.

### Reward sensitivity

In humans, the Behavioral Inhibition System (BIS) and the Behavioral Activation System (BAS) scales allowed us to approach behavioral motivation^58^.

In mice, reward sensitivity was evaluated at the end of the mGT using the sucrose preference test (adapted from Lutz *et al*.’s work^59^). For more details, see Supplementary data.

## Data analysis

### Whole group analyses

Population’s overall performances in terms of advantageous choices for the gambling tasks were divided in 5 blocks, each representing 20% of the task. For each block, the performances were compared to chance level using t-tests.

The evolution of performances was assessed by ANOVAs, with factor being 20%-Blocks. Differences between populations were analyzed by a partially repeated ANOVA with Species as the between subject variable and 20%-Blocks as the within subject variable.

The evolution of the choice behaviors’ scores were first compared for each group using a t-test. Then, scores from the overall populations were compared by two way repeated measures ANOVA with factors being Species and Time Course (first and last 40% of the task).

BIS/BAS data in humans and sucrose preference in mice were analyzed using Kruskal-Wallis (KW) tests (see Supplementary data).

Correlations between endpoint performances (percentage of advantageous choices in the last 30% of the task) and choice behaviors were also carried out.

Comparisons were Bonferroni corrected when necessary to account for multiple comparisons.

### Interindividual data analysis

For each group, the evolution of gambling performances was assessed by repeated measures ANOVAs, with factors being 20%-Blocks and Clusters (*good, intermediate* and *poor DMs*), followed by MW tests to further show subgroup differences two by two. Each interindividual distribution was also compared between mice and humans by using a MW tests.

We used W tests to compare, for each subgroup, the evolution of gambling performances from chance level. Subgroup’s proportions of each population were compared using a Chi-Square test.

Differences in choice behaviors’ scores between subgroups in mice and humans were assessed by KW tests, for the beginning and the end of the task. These measures were also compared between species using MW tests.

All MW and W tests were Bonferroni corrected.

The statistical significance threshold of all tests was set at p < 0.05.

## Acknowledgements

The authors thank Drs. Ruud van den Bos and Sylvie Granon for valuable advice to set up the mGT. We are also grateful to Mr. Hervé Reyssie from the *Animal Facilities, Besançon*, for technical support. This study was supported by grants from the Communauté d’Agglomération du Grand Besançon (LC), the French Eastern Interregional Group of Clinical Research and Innovation (GIRCI Est; appel à projet « jeunes chercheurs », APJ 2015; JG), which had no role in the study design, collection, analysis or interpretation of the data, writing in the manuscript, or the decision to submit the paper for publication.

## Author information

### Affiliations

Université Bourgogne - Franche-Comté, Laboratoire de Neurosciences Intégratives et Cliniques - EA481, 19, rue Ambroise Paré, Besançon, France Cabeza Alvarez L., Chabin T., Ramadan B., Joucla C., Nicolier M., Pazart L., Haffen E., Fellmann D., Gabriel D. & Peterschmitt Y.

Hôpital Universitaire CHRU, Clinical Psychiatry, Besançon, France, Giustiniani J., Nicolier M. & Haffen E.

Hôpital Universitaire CHRU, CIC-1431, Besançon, France Giustiniani J., Nicolier M., Pazart L., Haffen E. & Gabriel D.

### Contributions

Conceived and designed the experiments: LC, JG, DG, YP Performed the experiments: LC, JG, TC, BR, CJ, DG Wrote the paper: LC, JG, MN, LP, EH, DF, DG, YP

### Corresponding author

Peterschmitt Y.

