## Supplementary data for "Modeling decision-making under uncertainty: a direct comparison study between human and mouse gambling data"

**Supplementary Figure 1. Correlations between endpoint performances and behavioral determinants of DM.**
Correlations between endpoint performances and rigidity, flexibility, lose-shift and win-stay choices at the beginning and at the end of the task for human (white circles) and mice (black circles) populations.

**Supplementary Figure 2. Evolution of choice behaviors during task progression.**
Progression of the rigidity (A), flexibility (B), lose-shift (C) and win-stay (D) choices for humans (white circles) and mice (black circles) good, intermediate and poor DMs subpopulations, between the beginning and the end of the experiment. Comparison by repeated measures ANOVA and post hoc t-tests show slight differences between selected subpopulations across species (beginning versus end of the task: ns, non significant; #, $p<0.05$; ##, $p<0.01$; ####, $p<0.0001$; humans versus mice: *, $p<0.05$; **, $p<0.01$; ****, $p<0.0001$).

<u>Behavioral determinants of DM: subgroups' comparison.</u>
In humans, no difference in rigidity was found between subgroups at the beginning of the experiment (KW, $p>0.05$). On the contrary, good DMs were found to be significantly less flexible than intermediate and poor DMs (KW, $p<0.01$; vs intermediate: $p<0.01$; vs poor: $p<0.05$). At the end, strong differences from both scores were observed (rigidity: $p<0.0001$; flexibility: $p=0.0001$), good DMs being significantly more rigid and less flexible than the other two subgroups (rigidity: vs intermediate: $p<0.0001$; vs poor: $p<0.01$; flexibility: vs intermediate: $p<0.0001$; vs poor: $p<0.05$). No differences for lose-shift choices between subgroups were found at the beginning of the experiment ($p>0.05$), but at the end ($p<0.001$). Good DMs were more prone to choose the same option after a loss than intermediate ($p<0.001$), but not than poor DMs ($p>0.05$). Regarding win-stay choices, significant differences between subgroups were found at the beginning ($p<0.01$) and the end of the task ($p<0.0001$), being good DMs more prone to continue choosing the same option after getting a reward than the other two subgroups (vs intermediate: $p<0.01$; vs poor: $p<0.05$).

In mice, no differences were found between subgroups for rigidity, nor the flexibility, at the beginning of the experiment (KW, p>0.05). However, like in humans, significant differences emerged later in the task (KW, rigidity: p<0.001; flexibility: p<0.01). In fact, good DMs were also more rigid (p<0.001) and less flexible (p<0.01) than poor DMs, but with no difference compared to the intermediate group (p>0.05). No difference for lose-shift and win-stay choices between subgroups was observed at the beginning of the mGT (p>0.05), but at the end of the task (lose-shift: p<0.05; win-stay: p<0.01). In the same way as humans, mice good DMs were more prone to choose the same option after a penalty than intermediate DMs (p<0.05). Nevertheless, no difference was found compared to poor DMs. At the same time, good DMs used to remain in the same arm after a reward more frequently than poor DMs (p<0.01), but no differences were found regarding intermediate DMs (p>0.05).

**Supplementary Figure 3**. **Sensitivity to reward and/or punishment, assessed by BIS/BAS scales in humans and sucrose preference task in mice, does not differ between subgroups.**

BIS/BAS scales

In humans, no significant differences in BAS and BIS scores were observed between subgroups (KW, p<0.05).

Sucrose preference test

The sucrose preference was calculated as the percentage of sucrose solution (2,5% sucrose solution: D(+)Saccharose, Carl Roth, Karlsruhe, Germany) consumption relating to the total consumption from a test lasting 4 consecutive nights. After a *habituation phase* or *forced sucrose phase*, animals were isolated in individual cages at night and had exclusive access to two identical bottles, one containing drinking water and the other, the sucrose solution. Every morning bottles were removed and weighted, and animals were replaced in their home cages. The position of the bottles was switched between consecutive sessions in order to avoid the effect of place preferences.

In mice, no significant differences in reward sensitivity were found between subgroups (KW, p>0.05), although an increasing tendency could be identified from poor to good DMs.

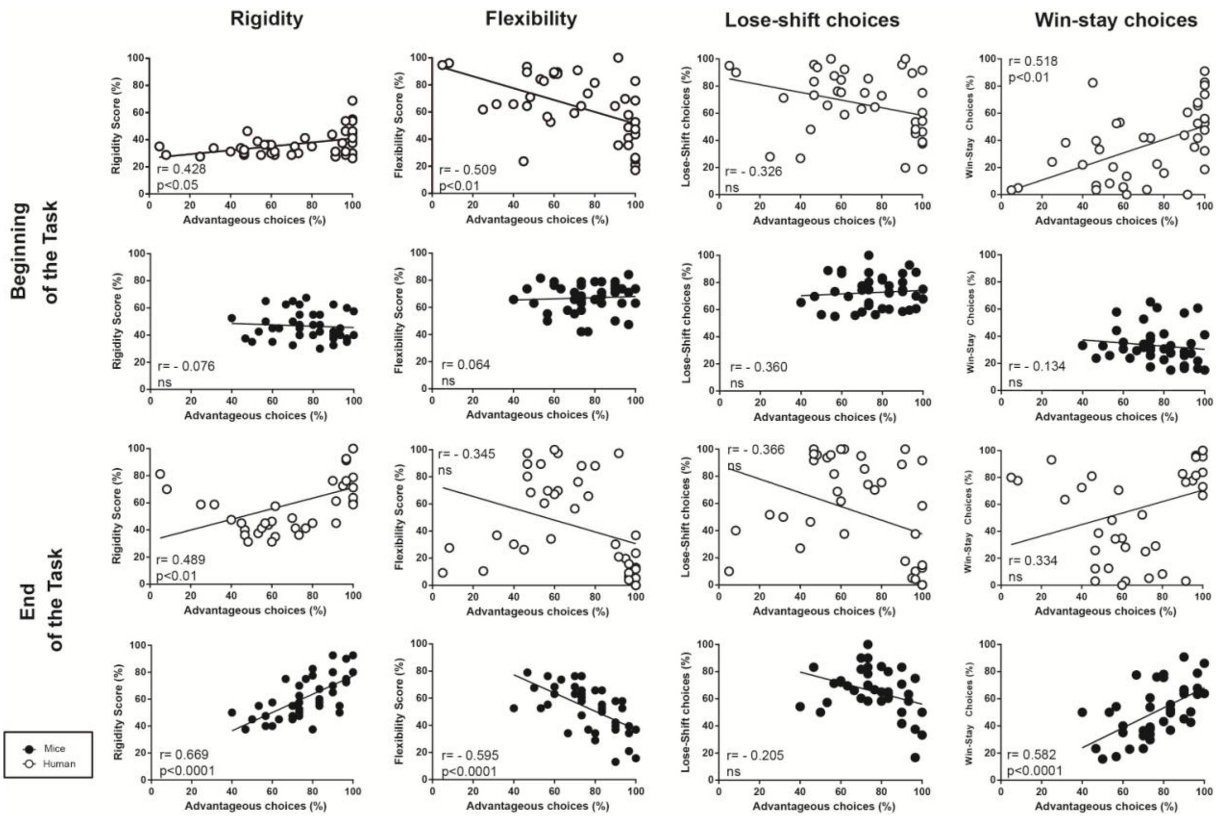

Rigidity
Flexibility
Lose-shift choices
Win-stay choices
Beginning of the Task
End of the Task
Rigidity Score (%)
Flexibility Score (%)
Lose-Shift choices (%)
Win-Stay Choices (%)
Advantageous choices (%)
r= 0.428
p<0.05
r= - 0.509
p<0.01
r= - 0.326
ns
r= 0.518
p<0.01
r= - 0.076
ns
r= 0.064
ns
r= - 0.360
ns
r= - 0.134
ns
r= 0.489
p<0.01
r= - 0.345
ns
r= - 0.366
ns
r= 0.334
ns
r= 0.669
p<0.0001
r= - 0.595
p<0.0001
r= - 0.205
ns
r= 0.582
p<0.0001
Mice
Human

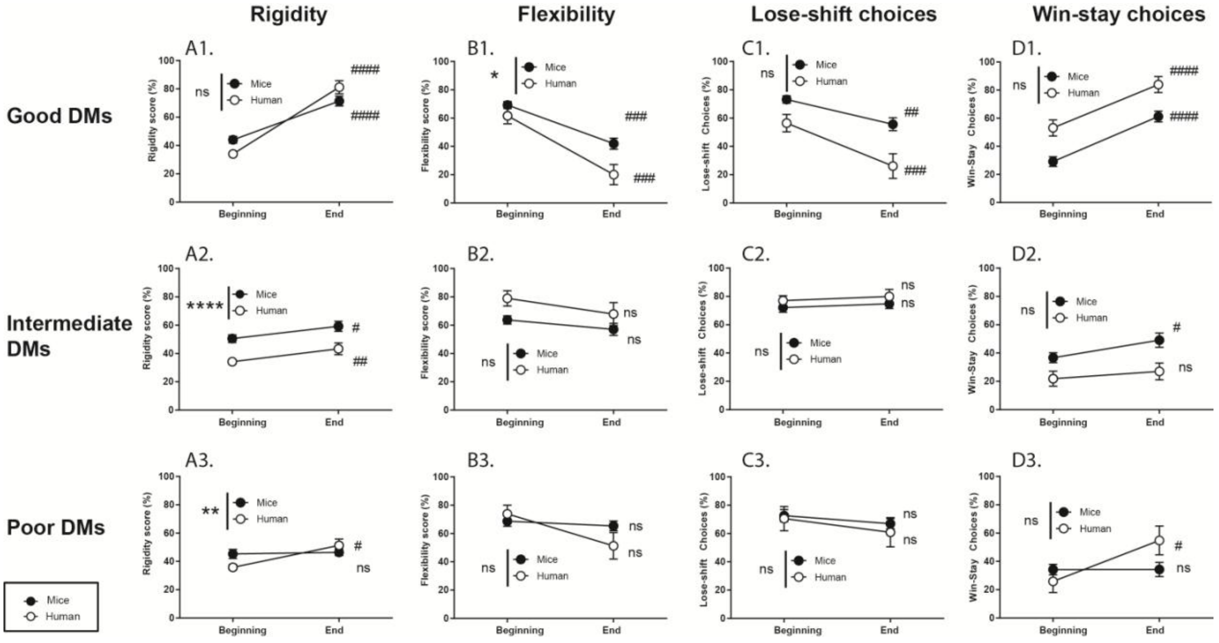

Rigidity
Flexibility
Lose-shift choices
Win-stay choices
Good DMs
Intermediate DMs
Poor DMs
A1.
B1.
C1.
D1.
A2.
B2.
C2.
D2.
A3.
B3.
C3.
D3.
Mice
Human
Beginning
End
Rigidity score (%)
Flexibility score (%)
Lose-shift Choices (%)
Win-Stay Choices (%)

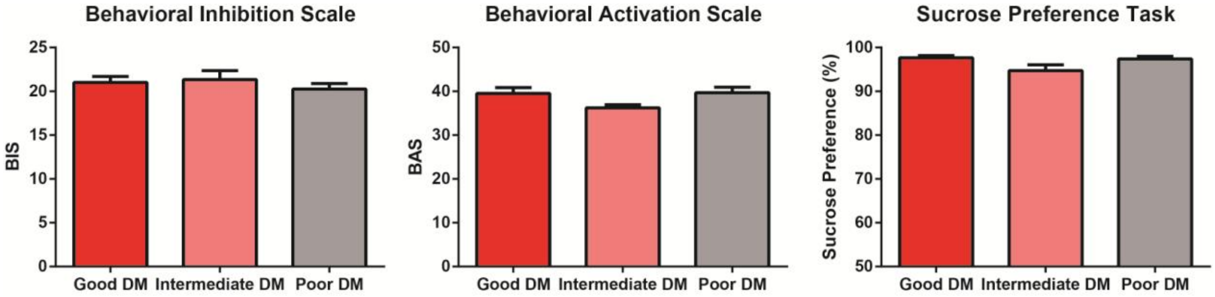
Behavioral Inhibition Scale
BIS
25
20
15
10
5
0
Good DM
Intermediate DM
Poor DM
Behavioral Activation Scale
BAS
50
40
30
20
10
0
Good DM
Intermediate DM
Poor DM
Sucrose Preference Task
Sucrose Preference (%)
100
90
80
70
60
50
Good DM
Intermediate DM
Poor DM
